# Beyond taxonomic identification: integration of ecological responses to a soil bacterial 16S rRNA gene database

**DOI:** 10.1101/843847

**Authors:** Briony A. Jones, Tim Goodall, Paul B.L. George, Soon Gweon, Jeremy Puissant, Daniel Read, Bridget A. Emmett, David A. Robinson, Davey L. Jones, Robert I. Griffiths

**Affiliations:** UK Centre for Ecology and Hydrology, Bangor, UK; UK Centre for Ecology and Hydrology, Wallingford, UK; School of Environment, Natural Resources and Geography, Bangor University, UK; School of Biological Sciences, University of Reading, RG6 6AS, UK

## Abstract

High-throughput sequencing 16S rRNA gene surveys have enabled new insights into the diversity of soil bacteria, and furthered understanding of the ecological drivers of abundances across landscapes. However, current analytical approaches are of limited use in formalising syntheses of the ecological attributes of taxa discovered, because derived taxonomic units are typically unique to individual studies and sequence identification databases only characterise taxonomy. To address this, we used sequences obtained from a large nationwide soil survey (GB Countryside Survey, henceforth “CS”) to create a comprehensive soil specific 16S reference database, with coupled ecological information derived from the survey metadata. Specifically, we modelled taxon responses to soil pH at the OTU level using hierarchical logistic regression (HOF) models, to provide information on putative landscape scale pH-abundance responses. We identify that most of the soil OTUs examined exhibit predictable abundance responses across soil pH gradients, though with the exception of known acidophilic lineages, the pH optima of OTU relative abundance was variable and could not be generalised by broad taxonomy. This highlights the need for tools and databases to predict ecological traits at finer taxonomic resolution. We further demonstrate the utility of the database by testing against geographically dispersed query 16S datasets; evaluating efficacy by quantifying matches, and accuracy in predicting pH responses of query sequences from a separate large soil survey. We found that the CS database provided good coverage of dominant taxa; and that the taxa indicating soil pH in a query dataset corresponded with the pH classifications of top matches in the CS database. Furthermore we were able to predict query dataset community structure, using predicted abundances of dominant taxa based on query soil pH data and the HOF models of matched CS database taxa. The database with associated HOF model outputs is released as an online portal for querying single sequences of interest (https://shiny-apps.ceh.ac.uk/ID-TaxER), and as a DADA2 database for use in bioinformatics pipelines. The further development of advanced informatics infrastructures incorporating modelled ecological attributes along with new functional genomic information will likely facilitate large scale exploration and prediction of soil microbial functional biodiversity under current and future environmental change scenarios.

## Introduction

Soil bacteria are highly diverse^**1, 2**^ and are significant contributors to soil functionality. Sequencing of 16S rRNA genes has enabled a wealth of new insights into the taxonomic diversity of soil prokaryotic communities, revealing the ecological controls on a vast diversity of yet to be cultured taxa with unknown functional potential^**3**^. However, despite thousands of studies across the globe, we are still some way from synthesising the new knowledge on the ecology of these novel organisms recovered through local and distributed soil surveillance. This is because there is currently no formalised way of retrieving ecological information on reference sequences which match user-discovered taxa (either clustered operational taxonomic units or amplicon sequence variants). Whilst we have a wealth of databases and tools for characterising the taxonomy of matched sequences^**4-6**^, databases do not include any associated ecological information on sequences matches. Whilst new software has recently become available that uses text mining to return some ecological data on matched sequences to NCBI, this information is currently limited to descriptions of sequence associated habitat^**7**^.

Synthesising relationships between soil amplicon abundances and environmental parameters is now necessary to progress ecological understanding of soil microbes beyond those few organisms that are readily cultivated. Determining microbial responses across environmental gradients can inform on the realised niche widths of discrete taxa, and may indicate the presence of shared functional traits across taxa^**8**^. This information is now urgently needed for microbes as we move into a period of increasing genomic data availability for uncultivated taxa. Coupling data on taxon responses across environmental gradients with functional trait information potentially allows a mechanistic and predictive understanding of both biodiversity and ecosystem level responses to environmental change. For example, a large body of theory exists describing how species responses to environmental change affects ecosystem functioning^**9-11**^. Here functional “response” groups are defined as species sharing a similar response to an environmental driver; and functional “effect” groups refer to species that have similar effects on one or more ecosystem processes. The degree of coupling between response and effect groups can then allow prediction of functional effects under change. For instance if certain phylogenetic groups of taxa decrease due to environmental change, and these taxa also represent an effect group (eg these taxa possess a unique functional gene) then we can expect the function to also decrease. Conversely with uncoupled effect groups (eg responsive taxa all possess a ubiquitous functional gene), the system is likely to be more functionally resistant to change^**11**^. Applying such concepts to microbial ecology is a realistic ambition given the extensive availability of amplicon datasets coupled to environmental information, and the increasing feasibility of uncultivated microbial genome assembly from metagenomes or single cell genomics^**12**^.

The fast evolution of microbial taxa coupled with potential horizontal gene transfer has led to assumptions that microbial diversity may be largely functionally redundant^**8**^. However we know from large-scale amplicon surveys that there are distinct differences in soil bacterial composition across environmental gradients, with soil pH frequently observed as a primary correlate^**13, 14**^. This implies that different microbial phylogenetic lineages possess adaptations conferring altered competitiveness in soils of different pH; paving the way for future studies into the genomic basis, and thereby elucidating specific genetic “response traits”. There is also evidence that many specific bacterial functional capacities such as methanogenesis (an “effect” trait) are phylogenetically conserved and therefore may be less redundant^**15**^. Determining the degree of functional redundancy in taxa which respond across soil pH gradients, will permit new insight into the microbial biodiversity mechanisms underpinning soil functionality and resilience to change. Since soil pH is largely predictable from geo-climatic^**16**^ and land use features^**17**^; prediction of the abundances of individual bacterial taxa under environmental change scenarios is likely to be feasible. The immediate challenge is therefore to establish predictive frameworks for many soil bacterial taxa, which can be populated with genomic information as it becomes available; to ultimately facilitate predictions of microbial functional distributions.

We believe that attempts to progress understanding of the ecological attributes of environmentally retrieved bacterial taxa can be streamlined immediately by making better use of the extensive amplicon datasets that exist, which already provide an abundance of useful information on taxa-environment responses. Indeed it has recently been shown that many prokaryotic taxa are distributed globally (particularly dominant OTUs^**18**^), yet there is currently no way to formally capture their ecological attributes in databases for further microbiological and ecological enquiry other than in supplementary material spreadsheets. Here we seek to address this by making available a database of representative sequences from a large 16S rRNA amplicon dataset from over 1000 soil samples collected across Britain. In addition to providing standard taxonomic annotation, we also seek to add ecological response information to each representative sequence. We focus here on soil pH responses as bacterial communities are known to respond strongly across soil pH gradients^**14**^. We will firstly model OTU abundances across soil pH using hierarchical logistic regression (HOF)^**19**^, a commonly used approach to examine vegetation responses across ecological gradients which has yet to be widely applied to microbial datasets. We will use model outputs to assign each OTU to a specific pH response group based on abundance optima, and in addition demonstrate the utility of the database in determining the phylogenetic relationships in ecological responses. The utility of the database will be further tested on 16S datasets to compare both the percentage of hits and modelled responses. The OTU database with associated HOF model outputs is released both as an online portal for visualising individual queries and as flat files for integration into existing bioinformatics pipelines.

## Results and discussion

### Database Coverage

The database was constructed from sequences obtained from the 2007 Countryside Survey (CS), a random stratified sampling of most soil types and habitats across Great Britain, full details of which are provided elsewhere^**14, 20**^. Sequencing of 1113 soils using the universal 341f/806r ^**21**^primers targeting the V3 and V4 regions of the 16S rRNA gene yielded a total of 39952 reference sequence OTUs, after clustering at 97% sequence similarity and singleton removal. Coverage was assessed on a filtered dataset of 1006 samples which had at least 5000 reads per sample, using sample based species accumulation curves calculated per habitat class and pooled across all habitats (**Fig.1**). The curves for individual habitats, whilst not reaching saturation, reveal some interesting trends with grasslands exhibiting highest biodiversity at the landscape scale, which is likely attributable to the broad range of soil conditions they encompass. The pooled curves across all habitats however appear to begin to level off, which importantly reveals that in total the reference sequence dataset provides good coverage of the non-singleton 97% OTUs found across this landscape.

### Performance of database against independent datasets

The coverage of this dataset was further assessed through blasting representative sequences from independent 16S datasets from various locations and habitats, against all 39952 CS representative sequences (**Table 1**).

**Table 1.** Validating the use of the CS OTU sequences as a database, through querying with independent datasets. Reference sequences from independent datasets were BLAST searched against countryside survey representative sequences, and the proportion of OTUs matched at over 97% similarity reported. British soil query datasets had highest percentage of hits irrespective of methodologies, with a set of riverine samples showing lowest proportion of OTUs matching the CS soil reference database.

| Query Dataset | Habitat Description | Query OTU percentage of hits | Primer | ENA Project ID |
| --- | --- | --- | --- | --- |
| 1 | Grassland and arable soils, Britain | 67.26% | 341f/806r V3-V4 | PRJEB36119 |
| 2 | All habitat soils survey, Wales | 58.49% | 515f/806rB V4 | PRJEB27883 |
| 3 | Thames River, Britain | 33.2% | 341f/806r V3-V4 | Unpublished, see Read et al, 2015 |

Here we defined an OTU ‘hit’ as a query OTU that shared 97% identity with a CS OTU and had an evalue equal to or less than 0.001. We subsequently calculated the percentage of OTUs within the independent dataset meeting this criteria to gain insights into coverage.

For the two soil datasets, we found over 50% of the OTUs in each study had a hit within the CS database. Expectedly, this was in stark contrast to a fresh water dataset which exhibited much less overlap with the CS database with 33.2% having CS hits. 16S sequences from dataset 1 (**Table 1**), a study of land use change across the UK^**22**^, also sequenced with the same 341f/806r primer set, had the highest percentage of hits against the CS representative sequences (67.26%). Wider assessment of our own unpublished datasets using the exact same methodologies yield percentages of hits of 62% and 56% for soils from UK calcareous grasslands and tropical rainforests respectively. A separate survey of Welsh soils^**23**^ was also queried against the CS database, which used the commonly used Earth Microbiome primer set exclusively targeting the V4 region (as opposed to V3 and V4 targeted region used for the CS dataset). This dataset had a percentage of hits of 58.49% providing evidence that datasets amplified with other primer sets can be matched to the CS database with only marginal loss of coverage.

We next wanted to explore possible reasons for obtaining less than 100% coverage from query soil datasets, given the good coverage of the CS reference sequence database evident from the rarefaction curve (**Fig.1**). We predicted this discrepancy was caused by rare OTUs being unique to specific studies, and tested this by classifying the UGRASS OTUs into 1000 discrete abundance based quantiles (1 being the most abundant quantile and 1000 being the least). Plotting the proportion of query OTUs which matched to the CS database by query OTU abundance class, confirmed that less abundant query OTUs had less matches to the CS database (**Fig.2**). This adds weight to arguments that much of the rare taxa detected through amplicon sequencing could be spurious artefacts of the PCR amplification process^**24**^. Regardless of these issues, the high proportion of hits for dominant taxa in the query dataset validates the use of the large CS dataset as a comprehensive reference database.

### Modelling OTU responses to soil pH

Since the majority of the 39952 reference OTUs obtained across all CS samples likely derive from rare taxa with intrinsically little value for predictive modelling (low within-sample abundance, and occurrence across samples), we opted to only model taxa-pH relationships for those taxa which occurred in at least 30 samples. These taxa were selected from a cleaned dataset of 1006 samples which had at least 5000 reads per sample. Further examination of the species accumulation by sample curves for the resulting 13781 OTUs, revealed saturation implying that this dataset had complete coverage of common OTUs, defined by being present in at least 30 samples across Britain. Huisman-Olff-Fresco models were then applied to determine individual bacterial taxa responses to pH using the R package eHOF using a poisson error distribution. Model choice was determined using AIC and bootstrapping methods implemented with the eHOF package ^**19**^, whereby the model with the lowest AIC was initially chosen and its robustness then tested by rerunning models on 100 bootstrapped datasets (created by resampling with replacement). If the most frequently chosen model in the bootstrap runs was different to the initial model choice, the most common bootstrap choice was selected. The resultant pH-taxa response curves classified by the HOF models include I: no significant change in abundance in response to pH, II: an increasing or decreasing trend, III: increasing or decreasing trend which plateaus, IV: Increase and decrease by same rate (unimodal) and V: Increase and decrease by different rates causing skew (**Fig.3**).

The proportion of OTUs assigned to each model is shown in **Table 2**, and reveals that most of the soil OTUs exhibited some trend with soil pH, and with the unimodal skewed model (V) being the most commonly fitted model type (45.76%). OTUs were then assigned to pH response groups based on the fitted pH optima. We classified OTUs demonstrating an acidic preference if the fitted optima was below pH 5.2, based on previous data showing this represented a critical threshold for bacterial communities^**14**^, which was further confirmed by a similar regression tree analyses of this sequence dataset (not shown). This pH value also represents a critical threshold in microbial functioning^**25**^. Similarly, a second threshold was designated at pH 7, with OTUs exhibiting an optima above this being classed as neutral, and those between 5.2 and 7 classed as “mid”. Plateau model shapes (model III), were sometimes more difficult to classify, since two optima are provided which span the plateau, and in some cases these crossed the pH 5.2 and 7 thresholds. Whilst OTUs exhibiting this response were in the minority, we opted to assign a separate designation representing this range, for instance “acid to mid” for an OTU with two optima above and below pH 5.2. The proportion of taxa classified to each pH response group are shown in **Table 3**. This reveals that OTUs with acidic preference are in the minority, consistent with reduced bacterial biodiversity being frequently observed in acidic soils^**14**^.

**Table 2.** Percentage of 13781 CS OTUs fitted to each HOF model. Each OTU was classified to one of five HOF model types according to fitted relationships with soil pH. The different model response shapes are shown in Fig 3.

| Model fit | Percentage of Countryside survey OTUs |
| --- | --- |
| **V** (Skewed Unimodal) | 45.76% |
| **III** (Plateau) | 24.13% |
| **IV** (Unimodal) | 23.52% |
| **II** (Monotonic) | 6.11% |
| **I** (No trend) | 0.49% |

**Table 3.**
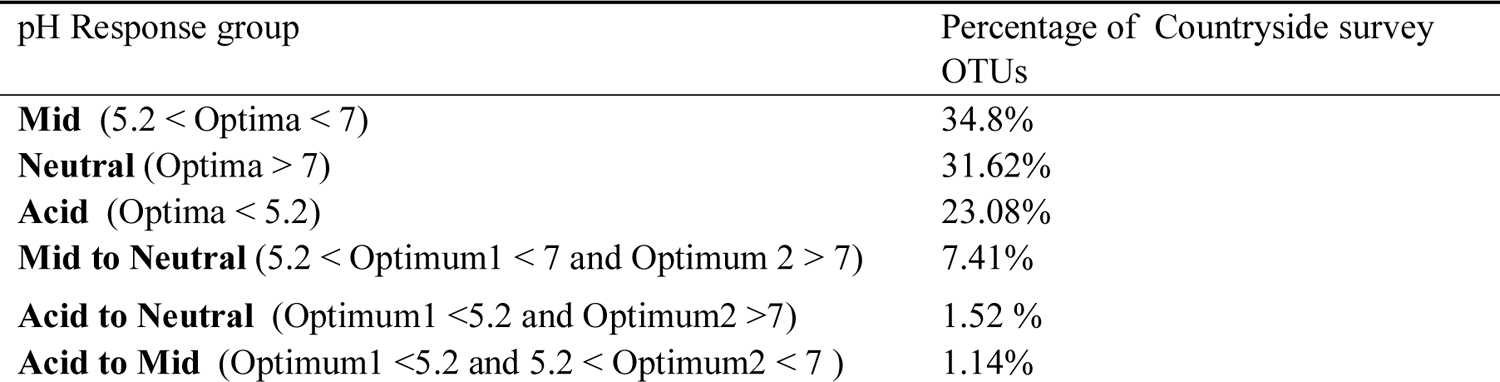
Percentage of 13781 CS OTUs classified to different pH response groups. Each OTU was assigned to a pH response classification based on the modelled pH optima. The model outputs with one optima (II, IV,V) were classified as acidic, mid or neutral based on pH thresholds identified above. Plateau shaped models with 2 optima (model III), which spanned the pH thresholds were labelled as either mid to neutral, acid to neutral, or acid to mid.

| pH Response group | Percentage of Countryside survey OTUs |
| --- | --- |
| **Mid** ($5.2 < \text{Optima} < 7$) | 34.8% |
| **Neutral** ($\text{Optima} > 7$) | 31.62% |
| **Acid** ($\text{Optima} < 5.2$) | 23.08% |
| **Mid to Neutral** ($5.2 < \text{Optimum1} < 7$ and $\text{Optimum2} > 7$) | 7.41% |
| **Acid to Neutral** ($\text{Optimum1} < 5.2$ and $\text{Optimum2} > 7$) | 1.52 % |
| **Acid to Mid** ($\text{Optimum1} < 5.2$ and $5.2 < \text{Optimum2} < 7$) | 1.14% |

Representative sequences of all 13781 OTUs were aligned with Clustal Omega 1.2.1 (http://www.clustal.org/), and used to construct a Phylogenetic tree with FastTree 2.1.7^**26**^ using neighbour-joining (NJ) with the generalized time-reversible (GTR) model of nucleotide evolution. The tree is shown in **Fig. 4** together with the pH classification derived from the HOF models. Distinct phylogenetic clustering is apparent for phyla with representatives known to have acidophilic preferences such as the Acidobacteria^**27**^. Additionally other phyla such as the Verrucomicrobia appear to possess clades with a distinct pH preference. However, the overall impression across other taxonomic groups is that the pH abundance optima can vary substantially amongst closely related taxa. This emphasises the need to move beyond the association of traits with broad phylogenetic lineages; and identifies the need to determine traits at finer levels of taxonomic resolution.

**Fig 1.**
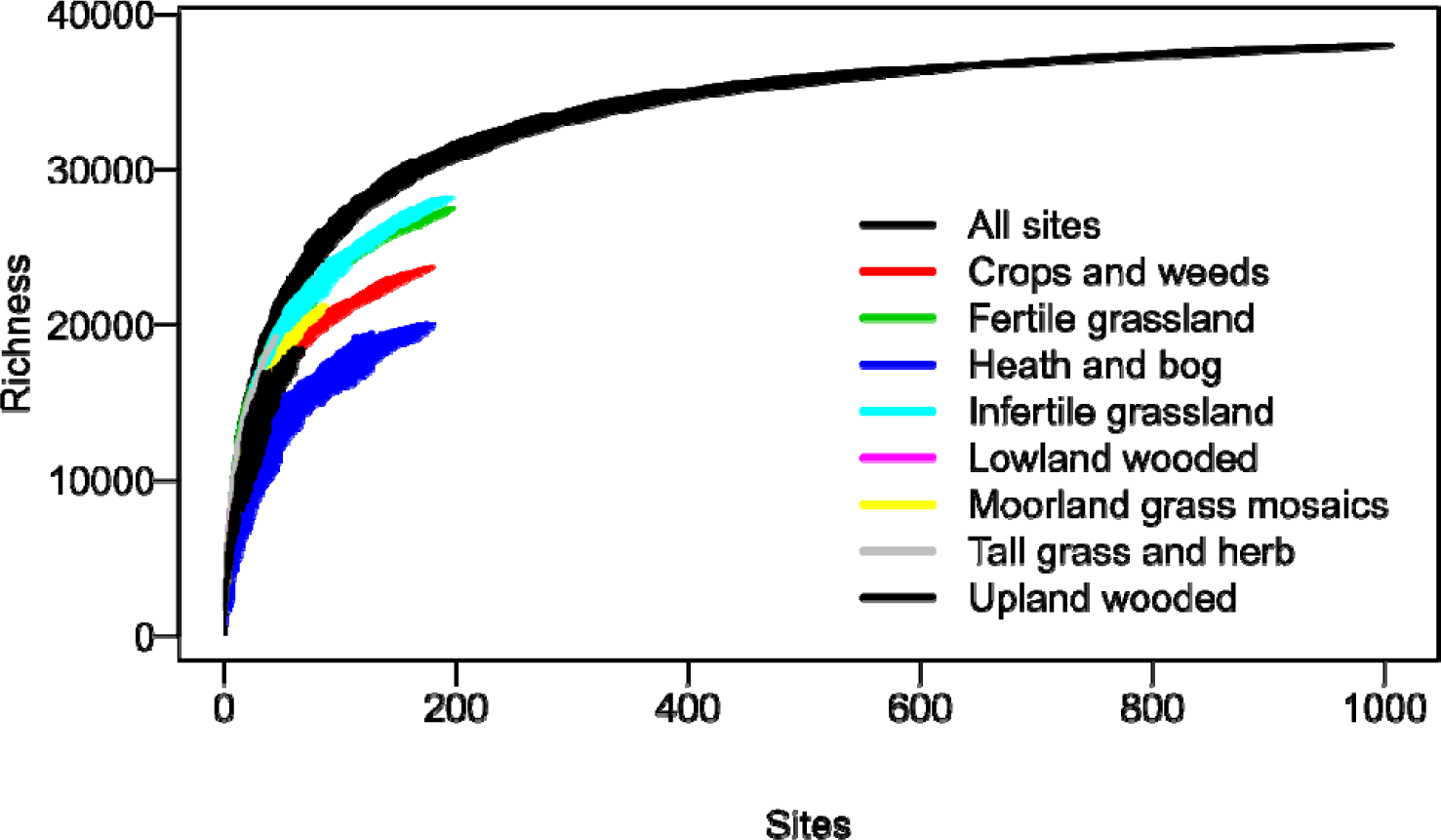
Coverage of bacterial 97% OTUs within the Countryside Survey (CS) dataset. Sample based richness accumulation curves were calculated across 1006 CS soil samples (“All sites”), and within specific habitats. Standard deviations are calculated from random permutations of the data.

**Fig 2.**
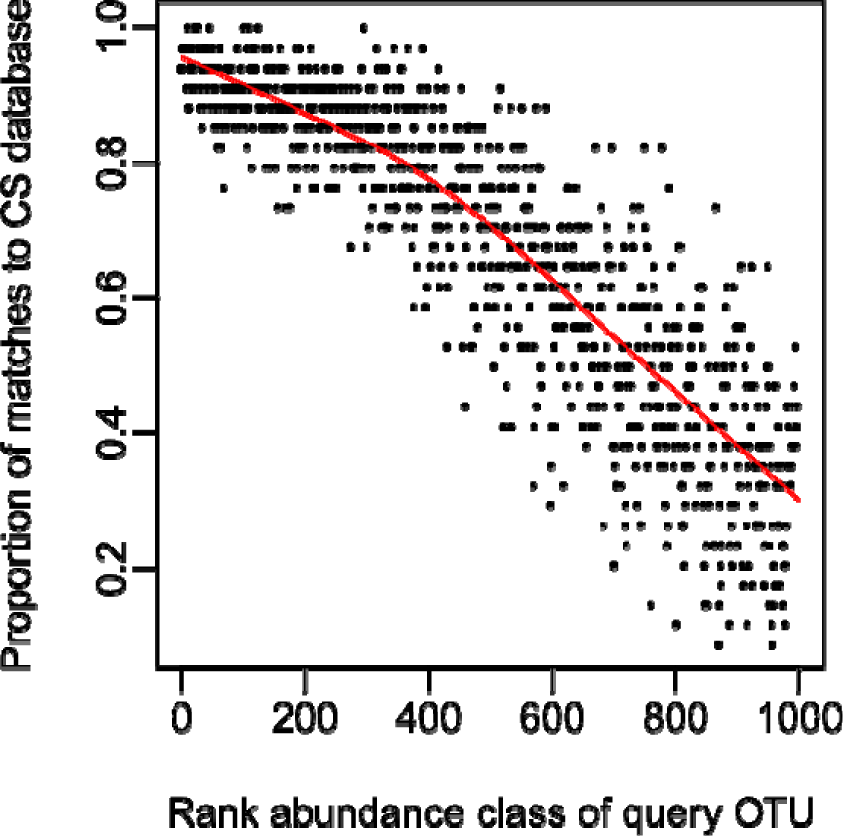
The CS database provides good coverage of dominant taxa within a query dataset. Query OTU reference sequences (dataset 1, table 1) were grouped into 1000 bins by decreasing rank (e.g the 1000th bin contains the least abundant OTUs); and the proportion of each bin matching the CS dataset calculated and displayed on the y axis. The proportion of matches to the CS database (> 97% similarity) declines as query taxa become rarer, despite the comprehensive nature of the CS database.

**Fig 3.**
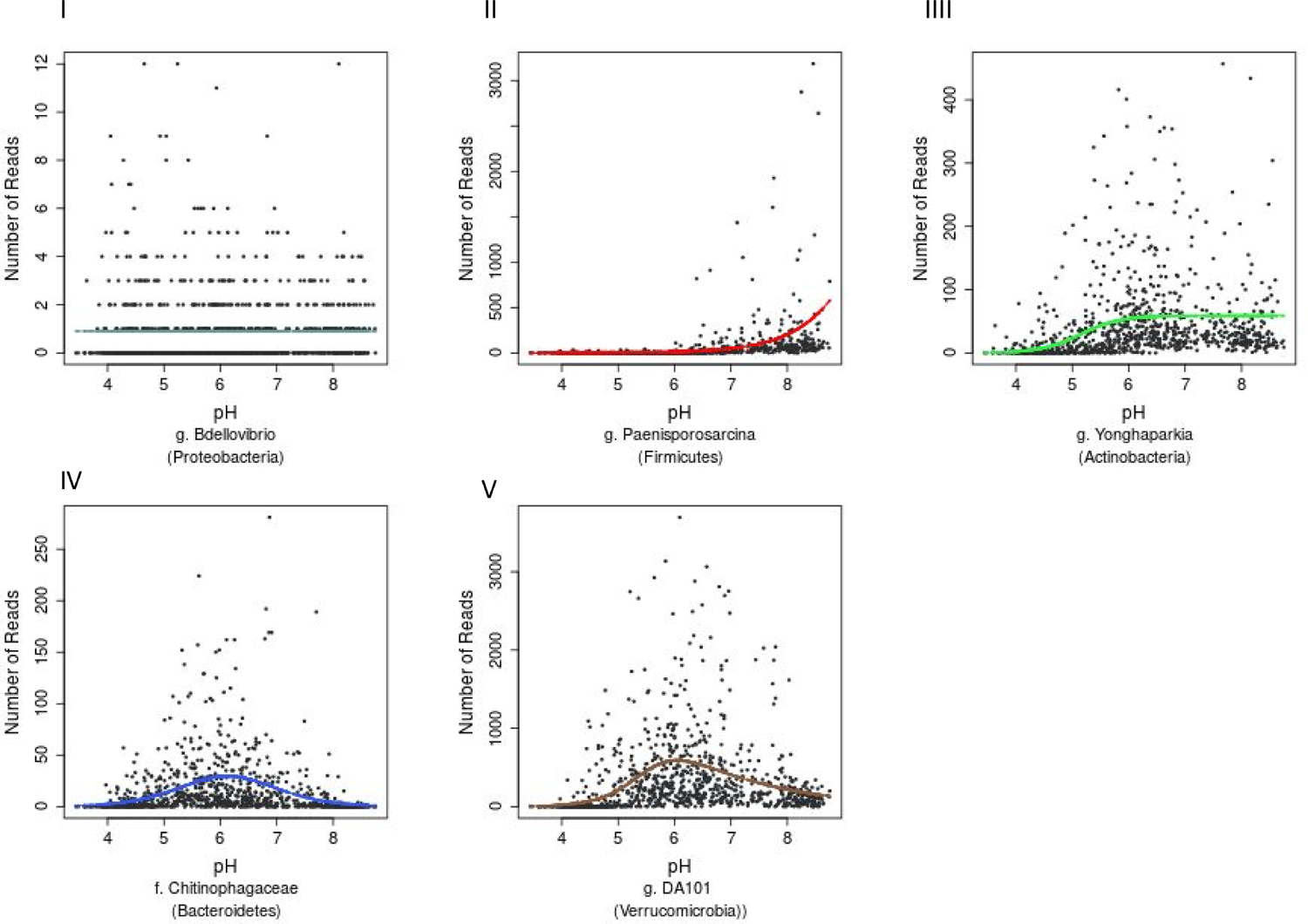
Examples of the five HOF model types. HOF models were generated through fitting countryside survey OTU abundances to soil pH (a pH range from 3.63 to 8.75). The five HOF models used were: I: no change in abundance across pH gradient, II: montonic an increase or decrease in abundance along pH gradient, III: plateau an increase or decrease in abundance along pH gradient that plateaus, IV: symmetrical unimodal, abundance increases and decreases across gradient at an equal rate, V: skewed unimodal, abundance increases and decreases across gradient at unequal rates.

**Fig 4.**
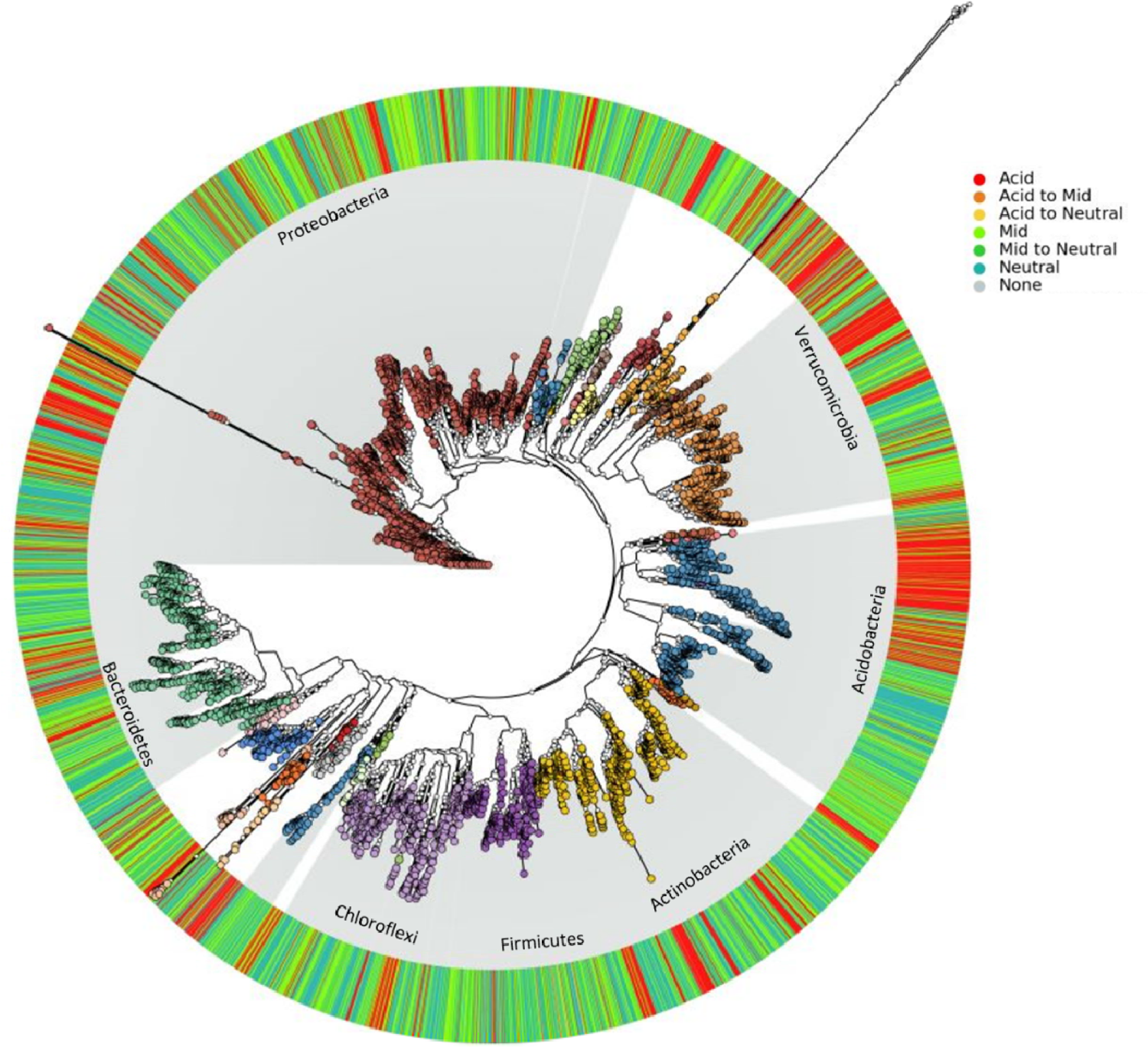
The phylogenetic distribution of bacterial pH optima. A phylogenetic tree of all OTUs present in >100 samples (totalling 6385 OTUs), with each OTU annotated according to pH classification based on HOF model optima (outer ring).

### Incorporating CS data and pH responses into a sequence identification tool

A web application was developed using the Shiny package (https://shiny.rstudio.com/) which enables users to BLAST a 16S query sequence against the countryside survey representative sequences, subsequently allowing visualization of key environmental information including HOF model outputs, relevant to individual matched sequences. The Graphic User Interface was implemented in R (3.4.1) using the Shiny package (https://shiny.rstudio.com/) alongside ShinyJS to execute JavaScript functions from R (https://cran.r-project.org/web/packages/shinyjs/). BLASTn commands are executed from R using the users query sequence, e value of 0.01, and the reference sequence database of CS representative sequences. eHOF model objects were converted to binary using the Rbase serialize function and stored in a PostgreSQL (9.3.17) database (https://www.postgresql.org/) alongside model and other environmental metadata (**Supp.fig.1**). BLAST results are displayed as an interactive table of hits, each hit linking to a plot of the pH model fit (based upon raw read number), a LOESS fit (based on relative abundance), a box plot of habitat associations and a simple interpolated map showing relative abundance distribution across Britain (**Supp.fig.2**). Additionally we provide a text box which can be populated with user submitted trait related information on matched OTUs. The application is available at shiny-apps.ceh.ac.uk/ID-TaxER/ and to facilitate batch processing of query sequences the sequence database, taxonomy and trait matrix are released via github (github.com/brijon/ID-TaxER-flat-files) for integration into bioinformatics pipelines.

### Utility in predicting pH preferences and community structure using a query dataset

To demonstrate both the utility of the reference sequence database, and the HOF modelling approach to identify environmental responses of soil bacterial taxa, we used a query dataset of >400 samples collected across Britain (dataset 1, **Table 1**). Since this survey focussed on productive habitats (grassland and arable land uses), with only a few acidic samples, it was not appropriate to generate independent HOF models. Instead we classified the samples according to the same pH cutoff levels identified above (pH 5.2 and 7) and then determined pH responsive taxa using indicator species analyses^**28**^. As can be seen in **Fig.5a**, the pH groupings were clearly evident in the sample based ordination. Representative sequences from this dataset were then blasted against the CS database, and optimum pH and pH classification metrics retrieved from the top hit for subsequent comparison. In total 477 indicators for the three pH groupings were retrieved, of which 454 had a match greater than 97% similarity to the CS database. Of the 155 acidic indicator taxa identified in the query dataset, 129 (83%) were reliably classified as acidic OTUs based on matches to the CS database (**Fig 5b**), with 20 OTUs “incorrectly” classified as having a mid-pH optima. However the predicted optima of these OTUs was mainly below pH 6 and most lie very close to pH 5.2. Similarly for the 226 query taxa identified as indicating neutral soils, 203 (90%) had a neutral pH classification in the CS database, with 15 being incorrectly classed as mid, though the optima for these was between pH 6.5 and 7. Sixty-seven indicators of the query mid pH soils were obtained of which 64 (96%) had a mid pH classification based on match to the CS database. Overall this analyses shows that information on soil pH preferences from independent datasets can be reliably obtained using our approach.

**Fig 5.**
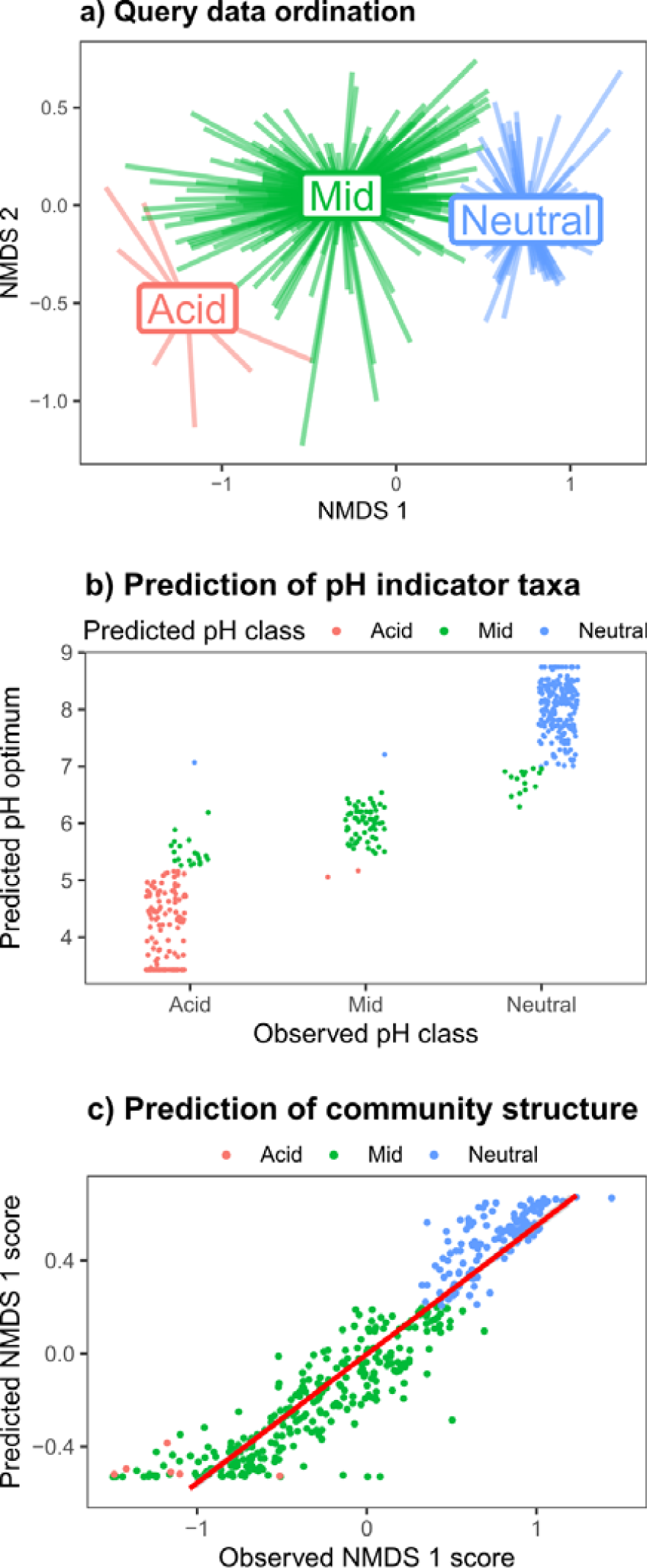
Validating the pH models using a query dataset. Taxa strongly responsive to soil pH were identified from Query dataset 1 (Table 1), and then matched to the CS database to evaluate utility of the approach. **a)** NMDS ordination plot of the query dataset, with pH groupings denoted by colour (red =pH<5.2; green=pH>5.2<7; and blue=ph>7). **b)** Indicator species analyses on the query dataset revealed 477 OTUs strongly associated with the three pH classes (“Observed pH class”). The y axis values and point colour denote the predicted pH optimum, and predicted pH class following matching to CS database. **c)** The relative abundances of the 100 most abundant taxa in the query dataset were predicted using the CS HOF models of matched taxa, and subjected to NMDS ordination. The plot shows that the predicted abundances of these taxa reliably predicted the observed data first axis NMDS scores.

We then sought to test whether we could reliably predict community structure using the CS HOF model outputs to predict query OTU abundances. We identified the most abundant OTUs in the query dataset, and blasted against the CS database. CS HOF models were then used to predict the abundances of the 100 matched dominant OTUs within the 424 query samples. This predicted community matrix was then subject to NMDS ordination with the first axis scores plotted against the actual observed ordination scores generated from 24260 OTUs. The results in **Fig 5c** show that the observed and predicted first axis ordination scores were highly related (*r*^2^ = 0.88) demonstrating that it is possible to predict broad scale community change from individual OTU relative abundance pH models. These findings add to a growing body of literature on the predictability of soil bacterial communities^**29-31**^; but furthermore demonstrate the utility of our overall approach in deriving meaningful ecological information from matches to a 16S rRNA sequence database incorporating ecological responses.

## Conclusions

This work demonstrates how large scale soil molecular survey data can be used to build robust predictive models of bacterial abundance responses across environmental gradients. The models were applied to the single soil variable of pH which is known globally to be the strongest predictor of soil bacterial community structure in surveys spanning wide environmental gradients. We have produced an informatics tool incorporating extensive sequence data from a wide range of soils, linked to taxonomic and ecological response information. This currently includes data on the modelled pH optima, and the predictive utility in this regard was demonstrated using an independent dataset. Other ecological information is also made available via an online portal including habitat association, spatial distribution, and metrics relating to abundance and occurrence. We are currently working on incorporating other information on the sensitivities of discrete OTUs to land use change; and there is the wider potential for users to update the trait matrix with other observations (more information provided at https://github.com/brijon/ID-TaxER-flat-files). Such information could include sensitivities to perturbations such as climate change, as well as rRNA derived links to wider genome data to inform on function.

We anticipate this simple database and tool will be of use to the soil molecular community, but also hope it prompts further global efforts to better capture relevant ecological information on newly discovered microbial taxa. We acknowledge some limitations of the current tool, and identify some possibilities to develop further: Firstly being a 16S rRNA amplicon dataset, the database inventory will be affected by known biases relating to PCR primers and amplification conditions^**32**^; and obviously, user datasets built on a different region of the 16S rRNA gene will not produce any matches. Additionally the length of sequences means only limited taxonomic resolution is currently provided, and ecological inferences based on BLAST matches must consider the strength of match, and variance within the matched region with respect to taxonomic discrimination^**33**^. Emerging long read sequencing technologies applied to survey nucleic acid archives in the future may improve these current constraints^**34**^. With respect to the pH models, many other factors can of course influence bacterial abundances^**3**, **35**^, and we note the large degree of variance in relative abundance for a taxon even within its apparent pH niche optima (**Fig 3**). Such variance could may be caused by nutrient availability, stress etc and more complex models, albeit constrained by pH, need to be formulated to advance predictive accuracy. More generally, we assert that observed taxon relative abundance only inform on relative taxon success at a given soil pH, and does not identify any explicit underpinning ecological mechanism (eg pH stress tolerance versus competitive fitness)^**36**^. However, linking emerging genomic data to detailed environmentally relevant sequence databases such as detailed here, will likely improve future understanding in relation to elucidating specific functional response traits and determining mechanisms underpinning bacterial community assembly along soil gradients. Finally, and importantly, the CS database is spatially constrained to a temperate island in Northern Europe, and would benefit from a more global extent to capture other soil biomes such as drylands. Improvements here could be made from integrating data from global sequencing initiatives, or leveraging data from sequence repositories provided consistent environmental metadata can also be retrieved in order to reliably predict response trait characteristics.

## Methods

Samples were collected as part of the UK Centre for Ecology and Hydrology Countryside survey (CS) between June and July 2007 covering sites throughout Great Britain. Samples were chosen through a stratified random sample of 1 km squares using a 15 km grid, implementing the institute of Terrestrial Ecology (ITE) land classification to ensure incorporation of different land classes, with up to 5 randomly sampled cores taken within each square. Metadata for each soil sample were collated including soil organic matter, soil organic carbon, bulk density, pH, indicator of phosphorus availability using methodologies detailed elsewhere^**14, 20**^.

DNA was extracted from 0.3g of soil using the MoBIO PowerSoil-htp 96 Well DNA Isolation kit (Carlsbad, CA) according to manufacturer protocols. Amplicon libraries were constructed according to the dual indexing strategy of Kozich et al^**37**^, using primers 341F^**38**^ and 806R^**39**^. Amplicons were generated using a high fidelity DNA polymerase (Q5 Taq, New England Biolabs) on 20 ng of template DNA employing an initial denaturation of 30 seconds at 95 °C, followed by (25 for 16S and 30 cycles for ITS and 18S) of 30 seconds at 95 °C, 30 seconds at 52 °C and 2 minutes at 72 °C. A final extension of 10 minutes at 72 °C was also included to complete the reaction. Amplicon sizes were determined using an Agilent 2200 TapeStation system (∼550bp) and libraries normalized using SequalPrep Normalization Plate Kit (Thermo Fisher Scientific). Library concentration was calculated using a SYBR green quantitative PCR (qPCR) assay with primers specific to the Illumina adapters (Kappa, Anachem). Libraries were sequenced at a concentration of 5.4 pM with a 0.6 pM addition of an Illumina generated PhiX control library. Sequencing runs, generating 2 x 300 bp, reads were performed on an Illumina MiSeq using V3 chemistry.

Sequenced paired-end reads were joined using PEAR^**40**^, quality filtered using FASTX tools (hannonlab.cshl.edu), length filtered with the minimum length of 300bp. The presence of PhiX and adapters were checked and removed with BBTools (jgi.doe.gov/data-and-tools/bbtools/), and chimeras were identified and removed with VSEARCH_UCHIME_REF^**41**^ using Greengenes Release 13_5 (at 97%). Singletons were removed and the resulting sequences were clustered into operational taxonomic units (OTUs) with VSEARCH_CLUSTER at 97% sequence identity. Representative sequences for each OTU were taxonomically assigned by RDP Classifier with the bootstrap threshold of 0.8 or greater using the Greengenes Release 13_5 (full) as the reference. All statistical analyses and visualisations were conducted within the R package, predominantly using the vegan and ggplot packages unless otherwise indicated.

## Acknowledgements

This work has been funded by the UK Natural Environment Research Council under the Soil Security Programme grant “U-GRASS” (NE/M017125/1) and an ENVISION DTP studentship award to Briony Jones.

---

**Supp. fig 1.**
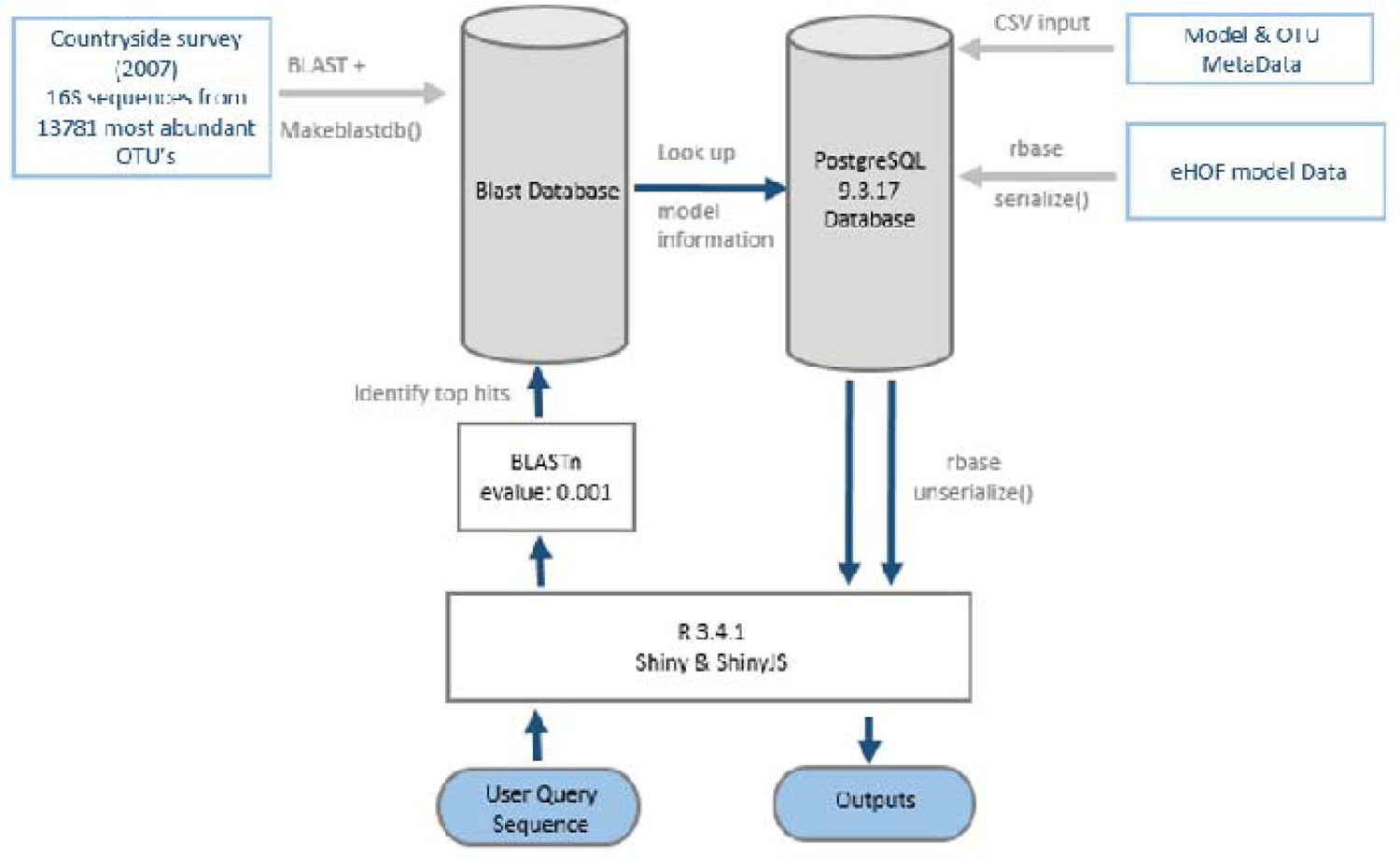
ID-TaxER database Infrastructure. 16S sequences are queried over the web via the R Shiny interface. A BLAST search is then performed against a blast database containing representative 16S sequences from the 2007 Countryside survey. Model information and associated metadata for match hits are located in a PostgreSQL database of OTU taxonomy/ model data, (model objects are stored as binary and retrieved for the user) and results displayed via the shiny interface.

**Supp. fig 2.**
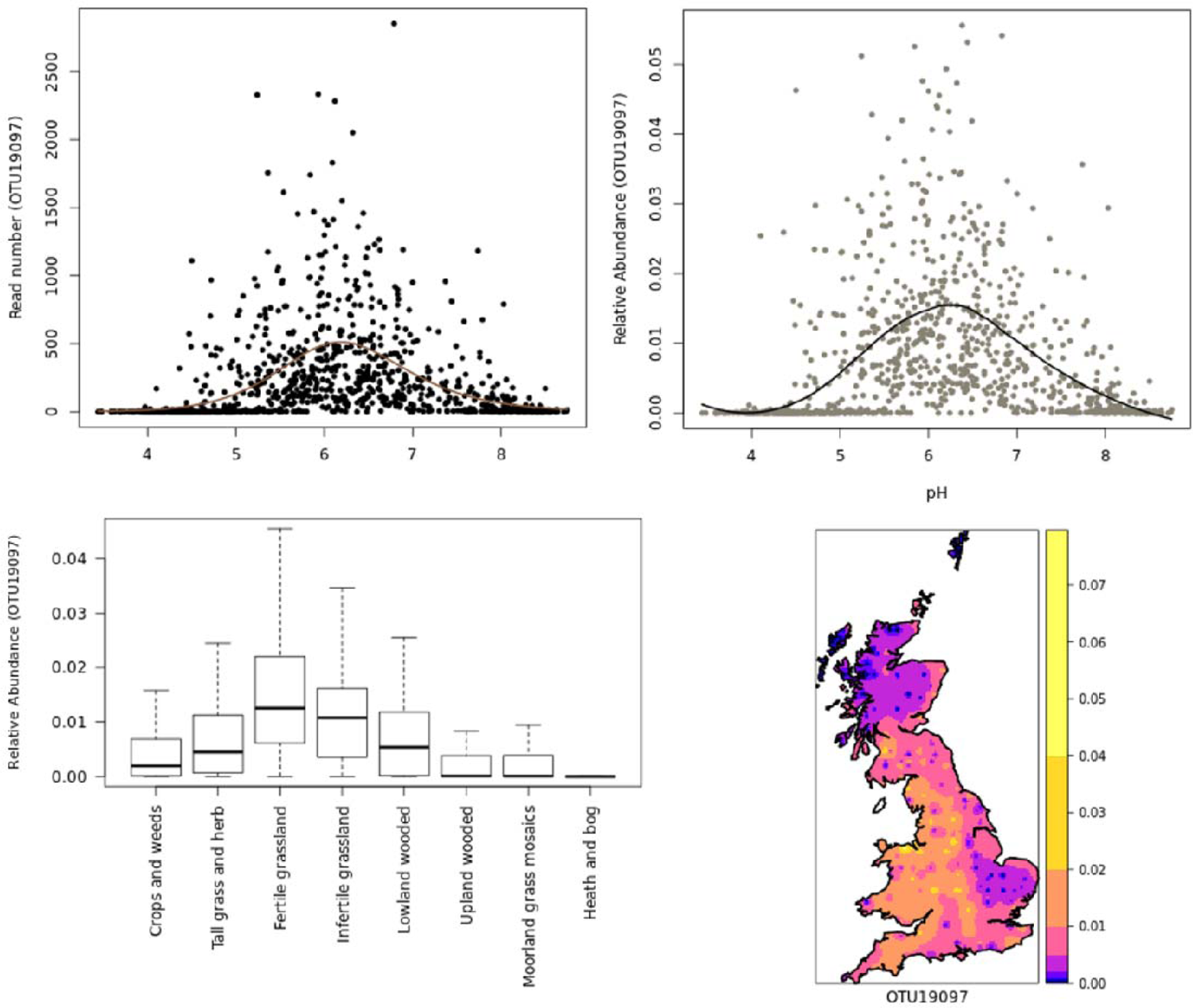
Example outputs from the ID-TaxER online portal. Using the DA101 /Ca. U. copiosus^**42**^ 16S sequence (GenBank: Y07576.1) as a query, we found 98.3% identitiy to CS OTU19097 (taxonomy=k_Bacteria; p_Verrucomicrobia; c_Spartobacteria; o_Chthoniobacterales; f_Chthoniobacteraceae; g_DA101): a) HOF model output showing the number of reads of CS OTU19097 per sample plotted against soil pH; with the line representing the model fit (Model V, unimodal response to pH with an optima at pH 6.18) b) the relative abundance of OTU19097 against sample pH, with the line representing a LOESS fit; c) boxplot showing the median and ranges of the relative abundance of OTU19097 per CS habitat class; d) inverse distance weighted interpolation map of the relative abundance of OTU19097 across Britain.

